# TUMOR-INFILTRATING NOCICEPTOR NEURONS PROMOTE IMMUNOSUPPRESSION

**DOI:** 10.1101/2024.08.23.609450

**Authors:** Anthony C. Restaino, Maryam Ahmadi, Amin Reza Nikpoor, Austin Walz, Mohammad Balood, Tuany Eichwald, Sebastien Talbot, Paola D. Vermeer

**Affiliations:** Cancer Biology and Immunotherapies Group, Sanford Research, Sioux Falls, USA; Department of Biomedical and Molecular Sciences, Queen’s University. Kingston. Canada; Department of Physiology and Pharmacology, Karolinska Institutet, Solna, Sweden

## Abstract

Nociceptor neurons impact tumor immunity. Removing nociceptor neurons reduced myeloid-derived suppressor cell (MDSCs) tumor infiltration in mouse models of head and neck carcinoma and melanoma. Carcinoma-released small extracellular vesicles (sEVs) attract nociceptive nerves to tumors. sEV-deficient tumors fail to develop in mice lacking nociceptor neurons. Exposure of dorsal root ganglia (DRG) neurons to cancer sEVs elevated expression of Substance P, IL-6 and injury-related neuronal markers while treatment with cancer sEVs and cytotoxic CD8 T-cells induced an immunosuppressive state (increased exhaustion ligands and cytokines). Cancer patient sEVs enhanced DRG responses to capsaicin, indicating increased nociceptor sensitivity. Conditioned media from DRG and cancer cell co-cultures promoted expression of MDSC markers in primary bone marrow cells while DRG conditioned media together with cancer sEVs induced checkpoint expression on T-cells. Our findings indicate that nociceptor neurons facilitate CD8+ T cell exhaustion and enhance MDSC infiltration. Targeting nociceptor-released IL-6 emerges as a novel strategy to disrupt harmful neuro-immune interactions in cancer and enhance anti-tumor immunity.

## Introduction

Head and neck squamous cell carcinomas (HNSCCs) are a collection of epithelial tumors that arise in the oral and oropharyngeal cavities (*1–3*); these cancers account for the sixth most common cancer diagnosed (*4*). HNSCCs are separated into two groups based on the mechanism of disease initiation; those induced by infection with high-risk human papillomaviruses (HPV^+^) and those that are mutationally driven through exposure to carcinogens such as alcohol and/or tobacco use (HPV^-^) (*5, 6*). While standard chemotherapy-radiation therapy provides excellent results for primary HNSCC, recurrence and metastasis remain a major challenge (*7, 8*). Recent advancements in immunotherapy offer additional options, but effects remain inconsistent with many patients failing to show improvement (*9*). As a result, efforts focused on identifying factors within the tumor environment that impact response to treatment continue. One such factor, includes the presence and functions of tumor-infiltrating nerves.

HNSCCs originate in areas rich with neuronal structures, including several cranial nerves. We have determined that HNSCC innervation primarily involves TRPV1-expressing nociceptor neurons from the trigeminal ganglia (TGM) (*10*). Similarly, melanoma is innervated by these neurons which are known to interact with both the innate and adaptive immune systems (*11, 12*). Neuro-immune crosstalk is crucial in regulating immune responses in infectious diseases (*13, 14*) and autoimmune disorders (*15, 16*). Recent studies in cancer models have also identified neural-immune interactions (*17*). In melanoma, sensory neurons facilitate the recruitment of myeloid-derived suppressor cells (MDSCs) and induce CD8^+^ T cell exhaustion via the release of calcitonin gene-related peptide (CGRP) (*18, 19*). However, the initiating mechanisms behind this neural-immune crosstalk in cancer are not fully understood.

Small extracellular vesicles (sEVs) are membrane-bound vesicles, measuring approximately 50-150nm in diameter, and are formed through the endocytic pathway (*20*). Released by all cells, sEVs serve as critical mediators of cell-to-cell communication both locally and at a distance (*21*). These vesicles carry a diverse array of biological materials, including proteins, lipids, and nucleic acids (*21*), and play a pivotal role in disease progression by contributing to metastasis and the remodeling of the tumor microenvironment (*22–24*). Integrins, embedded in the membranes of sEVs, are particularly crucial for targeting and preparing metastatic sites (*25*). The impact of sEVs on cancer progression, treatment resistance, and metastasis has been documented in various cancer, including HNSCC (*21, 26, 27*). Additionally, sEVs attract loco-regional neurons into the tumor milieu (*26, 27*), influencing local neuronal function and pain sensitization in murine models of oral cancer, thus underscoring their significant role in neuronal reprogramming and functionality (*27*).

We now employ syngeneic models of HNSCC and melanoma to explore the interaction between tumor-infiltrating nociceptor neurons, tumor cells and infiltrative immune cells. Our studies aim to determine the effects of nociceptor neuron loss on local tumor-associated immune cell populations. Additionally, we investigate whether tumors modify neuronal functions and, if so, assess how these changes contribute to the immunosuppressive environment within these cancers.

## Results

### sEV-mediated recruitment of nociceptor neurons is essential for disease initiation

The HPV^+^ HNSCC syngeneic murine cell line, mEERL cells, drive tumor innervation via release of sEVs (*26, 27*); this innervation promotes tumor growth as it is reduced when mEERL cells are implanted in nociceptor neuron-ablated mice (*10*). We have reproduced these data and now seek to examine whether compromising sEVs release in mEERL cells further impacts their growth in nociceptor ablated mice. To explore this, we used a mEERL cell variant where CRISPR-Cas9 technology deleted Rab27a and b, GTPases necessary for sEV release (*26*). When implanted into C57BL/6 mice, these cells exhibited reduced tumor growth (**Fig. 1**). Of note, tumor growth was completely blocked when the sEV-compromised cells (mEERL Rab27^-/-^) were implanted into nociceptor neuron-ablated mice (**Fig. 1**). This result was reproduced across three experiments involving 25 animals per group and suggests that sEV-mediated recruitment of nociceptor neurons is essential for disease initiation.

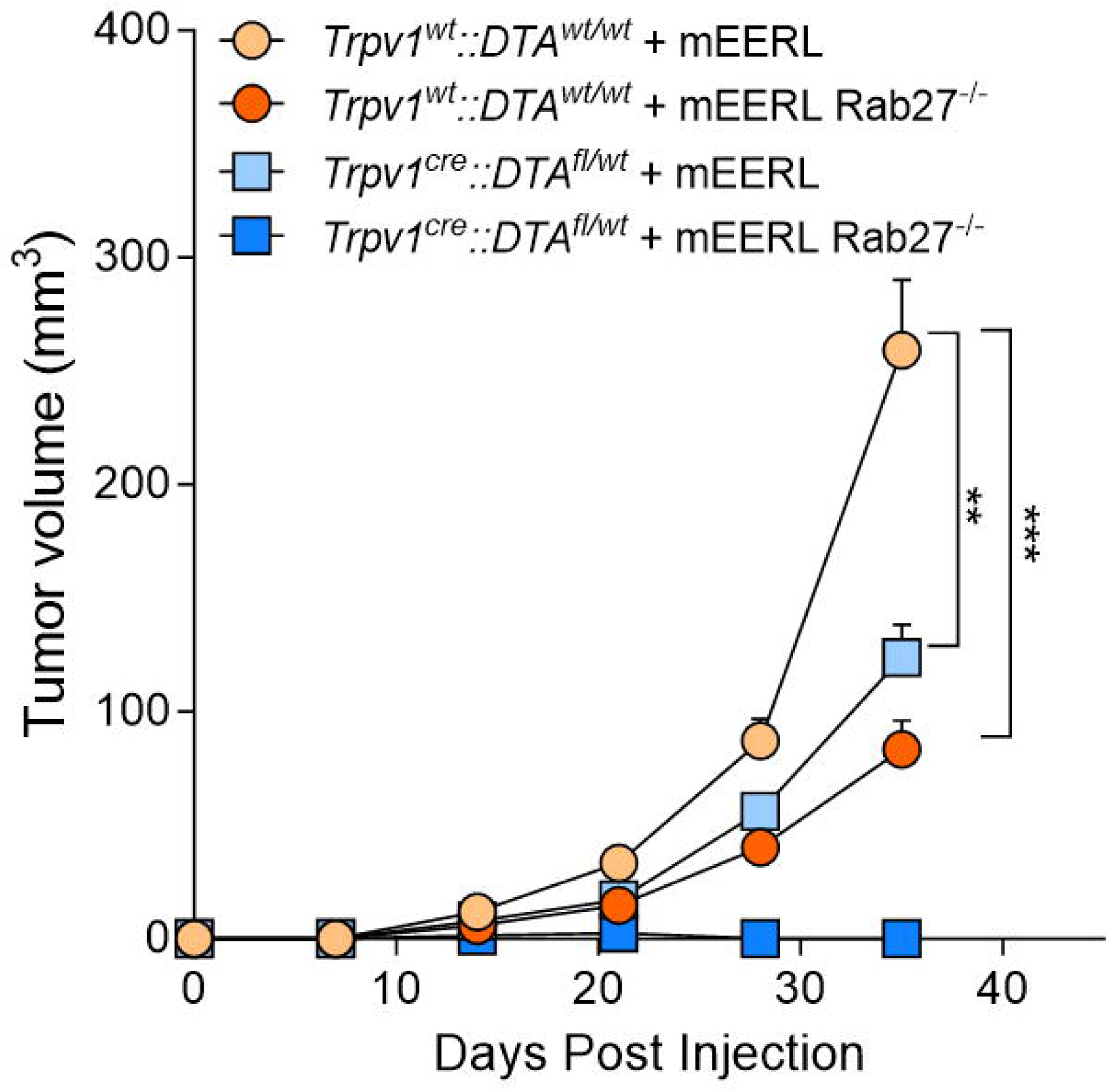

### Tumor-infiltrating neurons are transcriptionally modified

We hypothesized that tumor-released sEVs alter neuronal gene expression and function, thereby impacting disease initiation. To test this, mEERL cells were orthotopically implanted into C57BL/6 mice, and tumors were allowed to establish and grow. Twenty-five days post-tumor implantation, we harvested the ipsilateral TGM ganglia from tumor-bearing mice, with the contralateral TGM ganglia serving as controls. As expected from a previous study (*28*), the analysis of neuronal RNA from these ganglia revealed significantly increased expression of *Atf3*, a neuron-injury transcript, along with various regeneration-associated genes (RAGs) such as *Gap43*, *Gadd45*, and *Sppr1a* (**Fig. 2A**). Further validation with an additional group of mEERL tumor-bearing mice confirmed the increased expression of *Atf3* and markers of nociceptor neurons (*Cgrp*, *Tac1*) and *Tubb3*, a neuronal marker, in ipsilateral TGM neurons (**Fig. 2B**). Protein expression analysis by immunofluorescence confirmed our qPCR data (**Fig. 2C, D, Fig S1**).

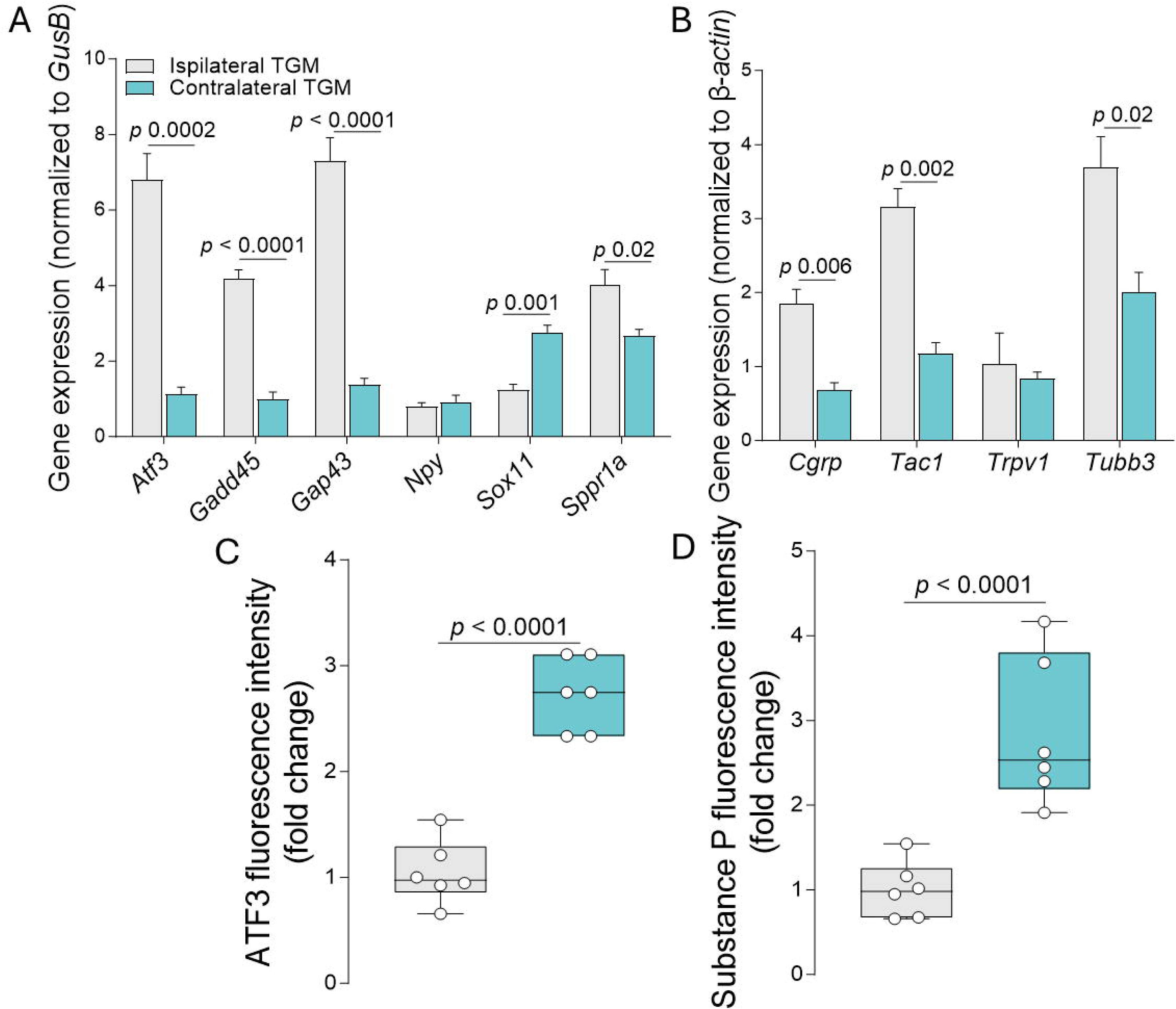

### Nociceptor neuron and mEERL cell interactions modulate the tumor milieu

This neuron injury gene signature in tumor-infiltrating neurons led us to explore whether these changes influenced the neuronal secretory profile. To investigate this, DRG neurons were co-cultured with mEERL cells, and the resulting conditioned media analyzed using a cytokine array. This co-culture resulted in an increased release of several factors, including IL-6, CCL2, CCL19, CXCL5, CD30L, CxCl16, TIMP1 (**Fig. S2**) and SP (**Fig. 3A**). In a neuronal co-culture with mEERL Rab27^-/-^ cells (compromised in sEV release), SP release was reduced (**Fig. 3A**). Further experiments tested whether SP directly induced IL-6 release from mEERL cells. While baseline IL-6 release was low in mEERL cells cultured alone, it significantly increased following treatment with recombinant SP. Notably, we had previously shown that mEERL cells express the SP receptor, NK1R (*10*); and now found that blocking this receptor in Substance P treated mEERL cells negated the IL-6 release (**Fig. 3B**). Additionally, DRG neurons themselves released more IL-6 when co-cultured with mEERL cells, an effect that was nullified by including an NK1R antagonist (**Fig. 3C**). As opposed to wildtype DRG neurons, we cultured DRG neurons from germline knockout IL-6 mice and found no detectable IL-6 in the conditioned media from these neurons when co-cultured with mEERL cells (**Fig. 3D, Fig. S2**). Il-6 release was increased when neurons were co-cultured with higher numbers of mEERL cells (**Fig. 3E**). Collectively, these findings suggest that interactions between mEERL cells and nociceptor neurons, mediated through both released sEVs and soluble factors like SP, led to increase IL-6 levels in the tumor microenvironment.

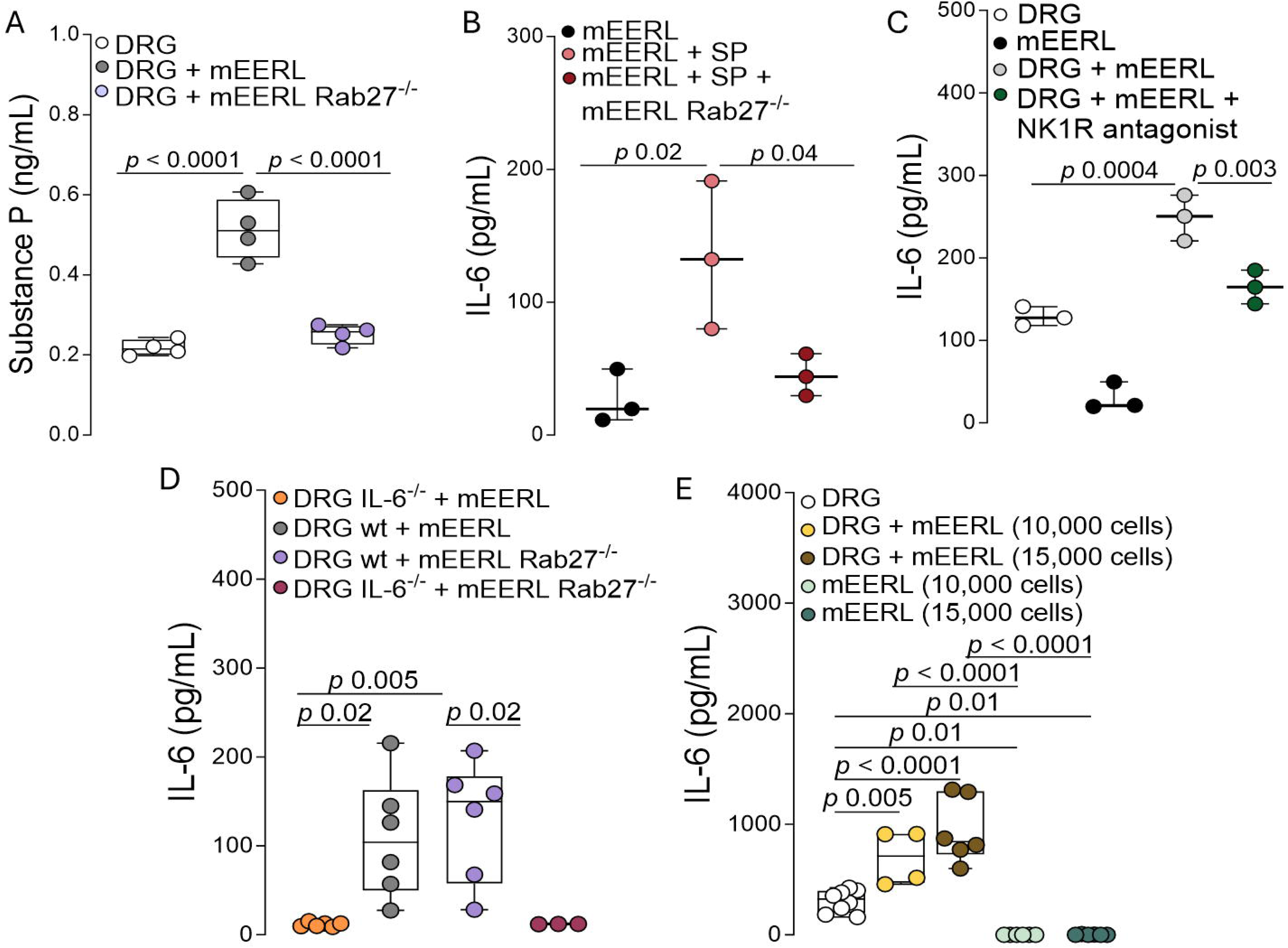

### Ablation of nociceptor neurons decreases MDSCs in the mEERL tumor bed

IL-6 has immunosuppressive action involving the expansion of MDSCs (*29–31*). Since we found that neurons are a major source of IL-6, we next sought to measure the impact of tumor-infiltrating nerves on tumor-infiltrating lymphocytes (TILs). We found that 25 days post tumor implantation, a phase of active growth and innervation, mEERL tumors implanted in nociceptor neuron ablated animals harbored a reduction in MDSCs and an increase in CD8^+^ T cells (**Fig. 4A**). Levels of CD4^+^ were not impacted (gating strategy shown in **Fig. S3**). To ensure that differences in tumor volume did not influence the tumoral MDSC population, we harvested tumors on day 15 post-implantation, when tumor volumes were comparable across the two groups. MDSCs are known to be a heterogeneous group of immune cells with two primary subtypes: monocytic MDSCs, which have high expression of Ly6C (Ly6C^hi^Ly6G^-^), and granulocytic MDSCs, marked by high expression of Ly6G (Ly6C^lo^Ly6G^+^) (*32, 33*). Despite their similar anti-tumor functions, these subtypes operate through distinctly different mechanisms (*34*). Detailed TIL immunophenotyping revealed that tumors from nociceptor neuron ablated mice had reduced numbers of granulocytic MDSC (**Fig. 4B**).

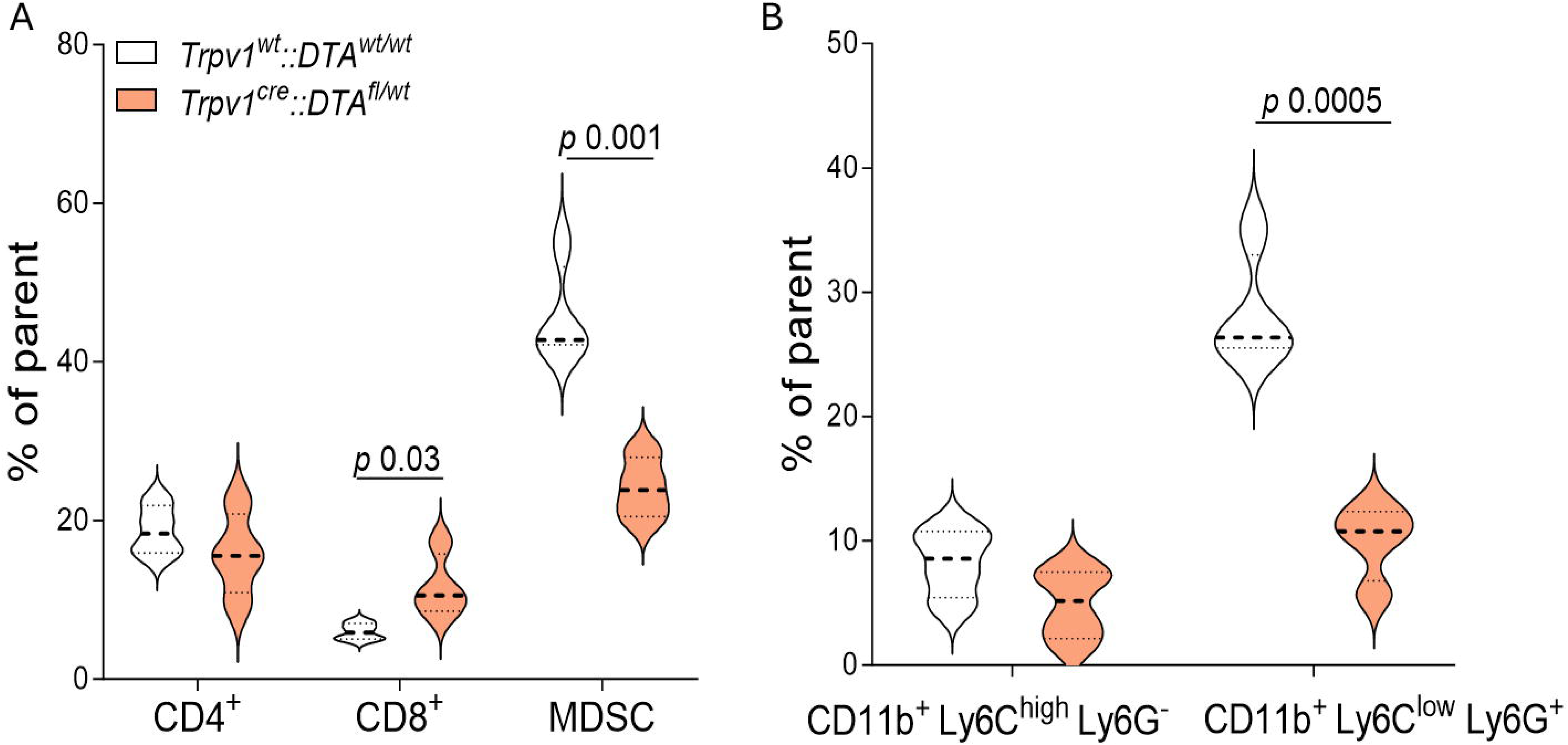

### Ablation of nociceptor neurons decreases MDSCs in melanoma and alters MDSC transcriptome

As tumor-infiltrating nociceptor neurons appear to drive this phenotype, we sought to test whether a similar phenotype could be observed in another densely innervated cancer. To test this, we used the syngeneic melanoma model B16F10-OVA cells implanted intradermally into either nociceptor ablated (*Trpv1^cre^::DTA^fl/wt^*) mice or their littermate controls (*Trpv1^wt^::DTA^fl/wt^*), the latter showing reduced tumor growth (**Fig. 5A**). Similar to the results seen in mEERL tumors, melanoma tumors from nociceptor neuron-ablated mice also showed a decreased MDSC population fourteen days post-tumor implantation (**Fig 5B**). Although the specific subgroups of MDSCs affected by the ablation of nociceptor neurons varied between mEERL and B16F10-OVA tumors, the consistent influence of these neurons on MDSC recruitment across different malignancies was evident. We next assessed the influence of nociceptor neurons on the MDCS transcriptome. To do so, we FACS-purified MDSC from nociceptor intact and ablated mice and profiled their transcriptome using RNA sequencing. We found ∼500 differentially expressed genes in nociceptor ablated mice, including decreases in *Csf1, Il17rb, Motch1, Tgfb1, Il10* and *Cxcl13* (**Fig. 5C, D**) all of which are known mediators of MDSC function (*35, 36*).

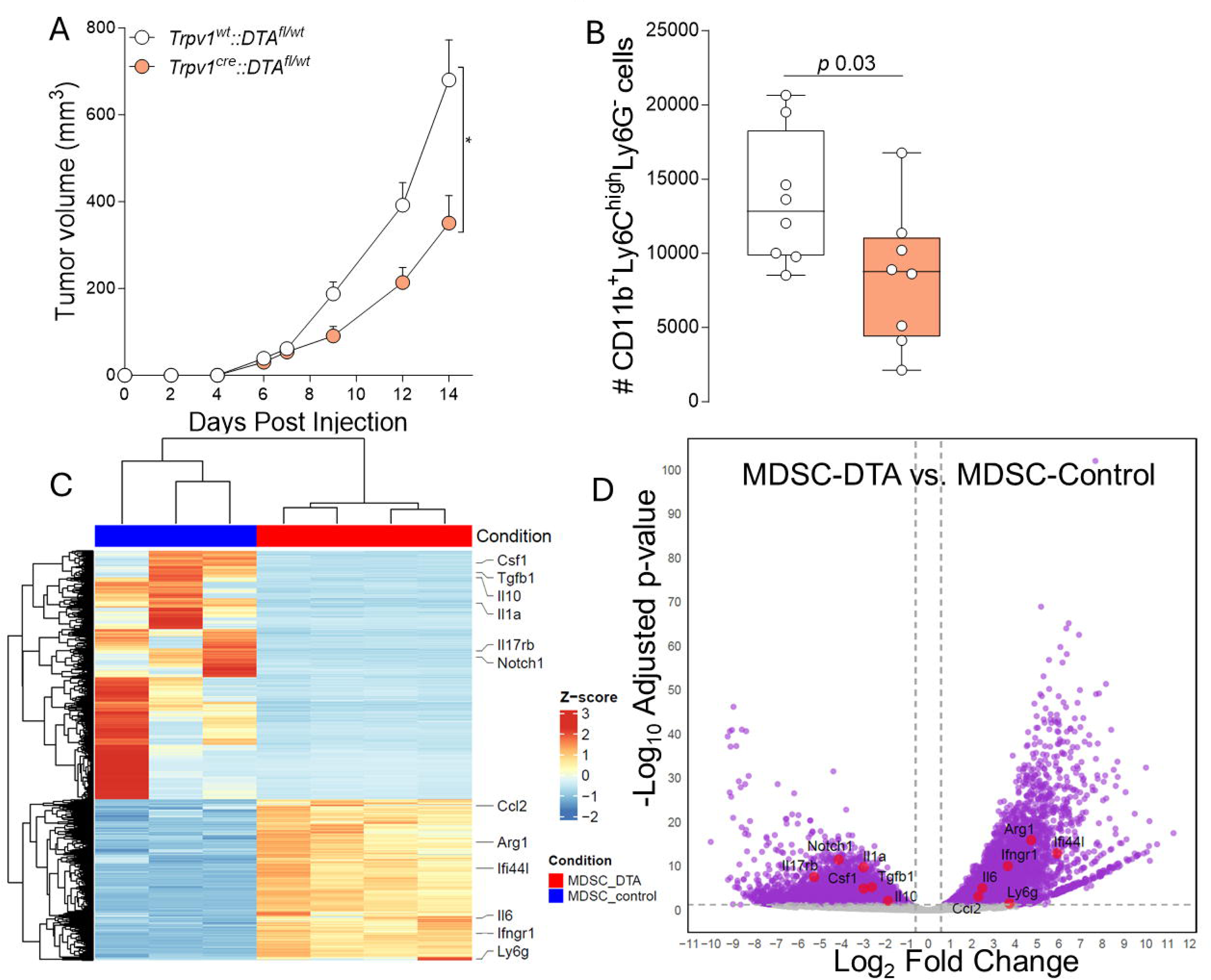

### Tumor cell-nociceptor neuron interactions induce MDSC differentiation from bone marrow cells

Given that nociceptor neurons modified the transcriptome of MDSC, and that IL-6 is known to drive their recruitment and expansion (*29*), we hypothesized that these tumor-infiltrating neurons might drive the differentiation of bone marrow cells into MDSCs. We tested this by co-culturing DRG neurons from nociceptor intact and ablated mice with sEV-competent or comprised mEERL cells and used the conditioned media to treat bone marrow cells harvested from C57BL/6 mice (**Fig. 6A**). Bone marrow cells stimulated with IL-6 and GM-CSF, which induces their differentiation into MDSCs (*37*), served as a positive control. Untreated bone marrow cells served as the negative control. As expected, IL-6 and GM-CSF stimulation significantly increased expression of CD11b^+^/Gr1^+^, MDSC markers, on the bone marrow cells. The induction of these markers was also robust in bone marrow cells stimulated with media from mEERL and wildtype DRG co-cultures but reduced with mEERL-Rab27^-/-^ cells and DRG co-cultures (**Fig. 6B**). These findings suggest that mEERL-released sEVs and DRG-released factors are crucial for MDSC differentiation. No significant effects on expression of these MDSC markers were noted with conditioned media that included nociceptor-ablated DRG, indicating a key role of nociceptor neurons (**Fig. 6B, gating strategy in Fig. S3**). Quantitative PCR analysis of treated cells showed that conditioned media from co-cultures of wildtype DRG and mEERL cells significantly induced *Arg1*, *Cox2*, and *Cybb* expression, essential for MDSC immune suppression (*38–40*) (**Fig. 6C**). These results highlight that mEERL cell/DRG interactions not only promote MDSC marker expression (CD11b, Gr1) but also genes vital for their immune functions. Other conditioned media had less impact, emphasizing the critical role of tumor cell/nociceptor neuron interactions in shaping the tumor microenvironment and immune cell phenotypes.

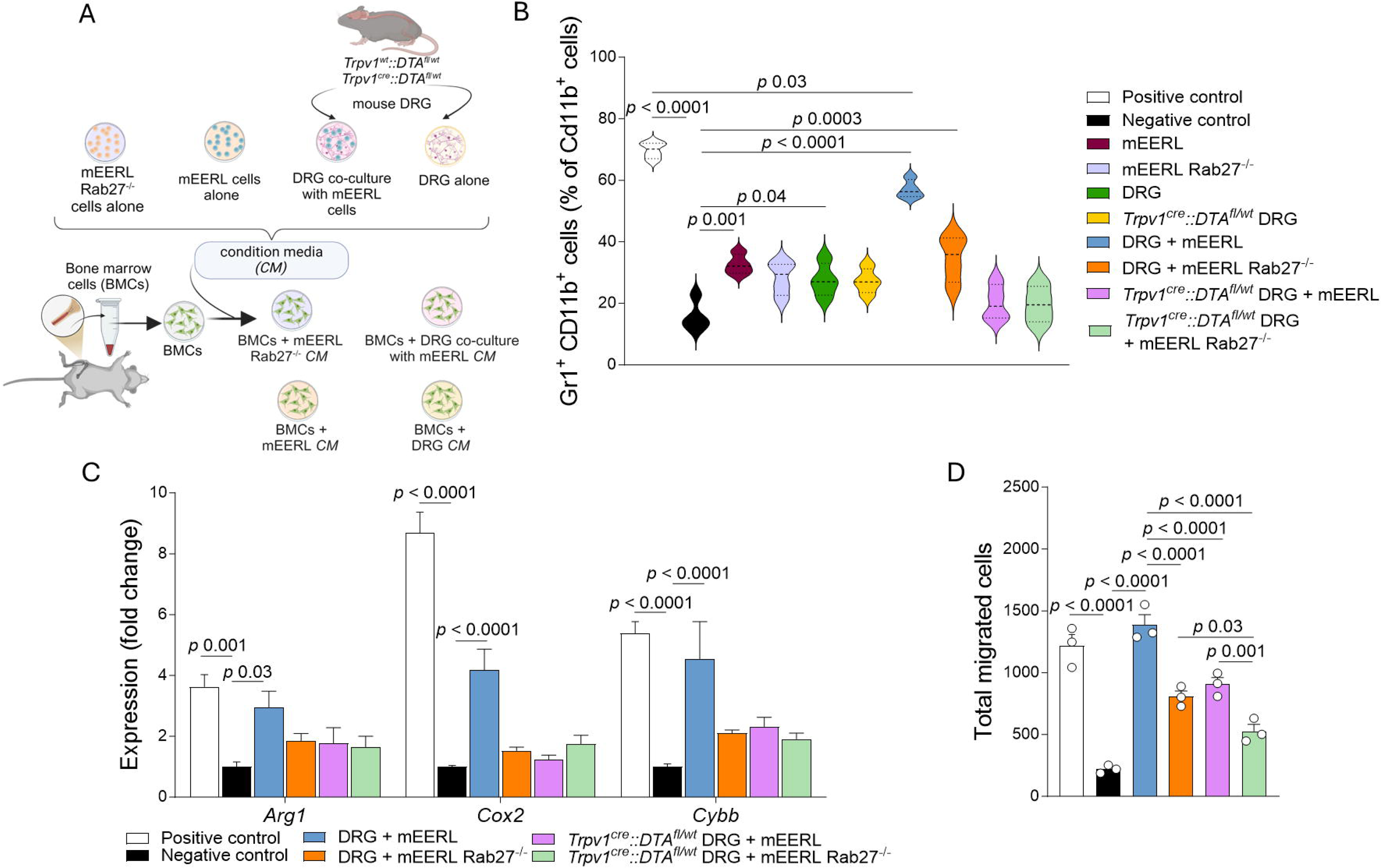

### Tumor cell-nociceptor neuron interactions induce MDSC migration

We next explored how mEERL/DRG interactions influence bone marrow cell migration to tumors. DRG from nociceptor intact and ablated mice were cultured with mEERL cells or their sEV-compromised variant (*26*). Concurrently, bone marrow cells from nociceptor intact mice were treated with GM-CSF and IL-6 to differentiate them into MDSCs expressing CD11b and Gr1 (*37*). After three days, these MDSCs were exposed to conditioned media from the mEERL/DRG co-cultures in a transwell assay. 24 h later, migration analysis showed that conditioned media from mEERL cells and wildtype DRG co-cultures led to the highest migration levels. In contrast, media from DRG co-cultured with Rab27^-/-^ cells, which have impaired sEV release, significantly reduced migration. These data highlight the strong migratory influence of mEERL-released sEVs. The migration levels from nociceptor neuron-ablated DRG with mEERL cells were similar to those from wildtype DRG with Rab27^-/-^ cells, indicating that nociceptor neuron-derived factors significantly boost MDSC migration. The lowest migration occurred with media from nociceptor-ablated DRG and mEERL-Rab27^-/-^ cells co-cultures (**Fig. 6D**).

### Nociceptor neurons and mEERL sEVs promote CD8^+^ T cell exhaustion

To this point, our data reveals that nociceptor neurons influence the tumor microenvironment by affecting both differentiation and recruitment of MDSCs to tumors, prompting further examination of their impact on other immune cells. Moreover, our data suggest that CD8^+^ T cells are increased in mEERL tumors from nociceptor-ablated animals (**Fig. 4A**). Notably, nociceptor interactions with CD8^+^ T cells during bacterial infections inhibit immunity by releasing neuropeptides (*13, 14*). Moreover, tumor released sEVs can directly impact CD8^+^ T cells (*41*). Thus, we investigated whether nociceptor neurons and/or tumor sEVs also modulate CD8^+^ T cells. Thus, we generated CD8^+^ T cells by activating splenocytes from C57BL/6 mice under T_c1_ inflammatory conditions. These CD8^+^ T cells were then exposed to conditioned media from wildtype DRG, mEERL sEVs, or both for four days. Analysis via flow cytometry showed that DRG media increased PD-1, LAG3, and TIM3 co-expression, and decreased IFNγ and IL-2 production. mEERL sEVs alone had no effect, but their combination with DRG conditioned media enhanced expression of these immune checkpoints and further reduced cytokine levels, indicating that nociceptor neuron-released factors and tumor sEVs jointly promote CD8^+^ T cell exhaustion (**Fig. 7A-C, gating strategy in Fig. S4**).

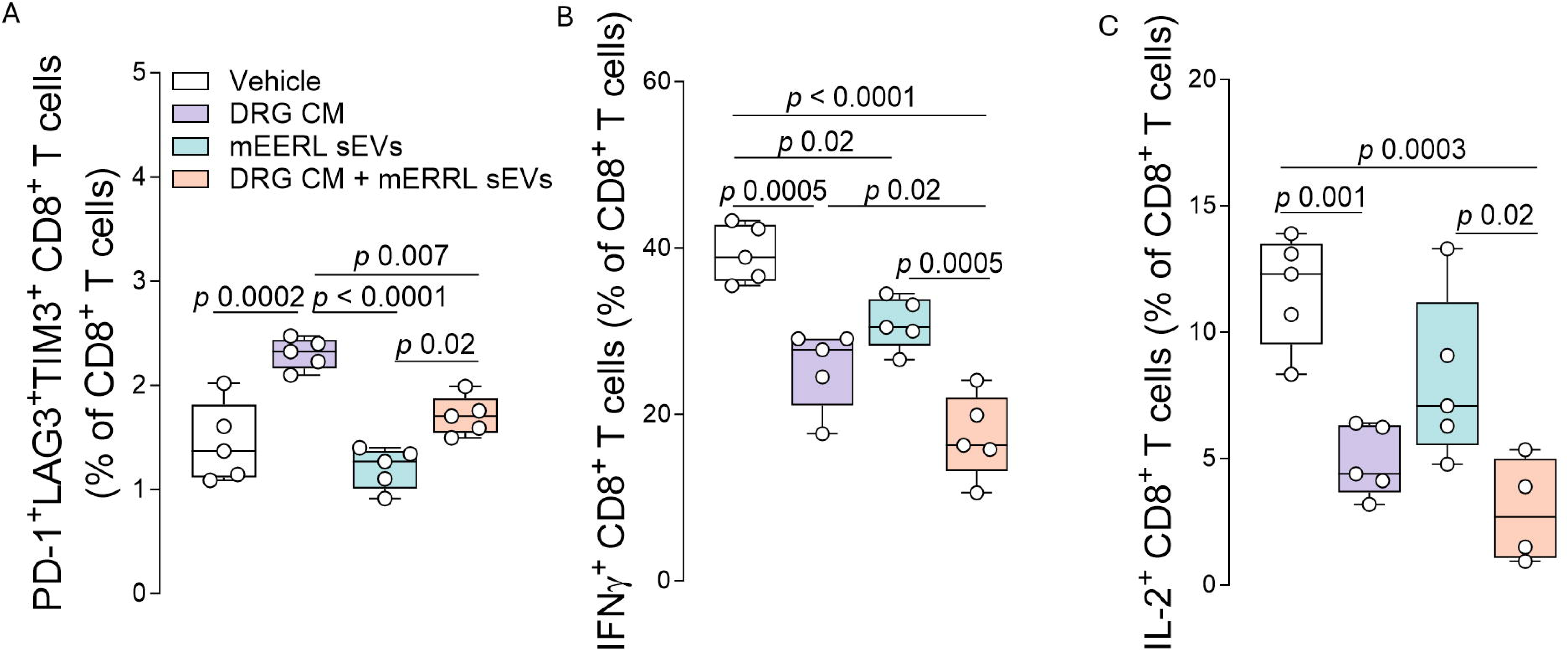

### mEERL sEVs and CD8^+^ T cells modulate the nociceptor neuron transcriptome

Our *in vivo* data (**Fig. 2**) demonstrate that tumor-infiltrating neurons are transcriptionally modified by the tumor microenvironment (TME). We hypothesized that mEERL sEVs or CD8^+^ T cells (or both) contribute this effect. To test this, DRG neurons from TRPV1^cre^::tdTomato^fl/wt^ reporter mice were cultured alone or with mEERL-derived sEVs and/or CD8^+^ T cells. After 48h, nociceptor neurons were FACS-purified, and RNA sequenced. Gene expression heatmaps indicate the influence of mEERL sEVs and CD8^+^ T cells on the transcriptome of nociceptor neurons. Among other changes, nociceptor neurons co-cultured with cytotoxic CD8^+^ T cells increased their expression of pro-exhaustion ligands (Pd1, also known as CD274) and cytokines (Il6), injury markers (Atf3, Sprr1a, Cryba2, Fgf3), and immunomodulatory neuropeptides (Gal, Calca; **Fig. 8A-B**). Interestingly, the exposure to mEERL sEVs had the opposite effect, decreasing some of the neuron reprogramming, including the injury markers (Atf3, Sprr1a, Cryba2, Fgf3; **Fig. 8C-D**). Finally, when combining both mEERL-derived sEVs and CD8^+^ T cells we drastically exacerbate nociceptor neurons reprogramming toward a pro-immunosuppressive phenotype; as shown by increased expression of pro-exhaustion ligands (Pd1) and cytokines (Il6), injury markers (Atf3, Sprr1a, Cryba2, Fgf3, Gpr151, Slc6a4, Ecel1), and immunomodulatory neuropeptides (Gal; **Fig. 8E-F**). Along with this transcriptomic reprogramming, we sought to test whether HNSCC patient-derived sEVs impact the response of nociceptor neurons to noxious ligands. As a proxy for this response, we examined the influx of calcium in responses to the TRPV1 agonist capsaicin (300 nM). Compared to neurons exposed to control sEVs, those exposed to sEVs from HNSCC patients showed an increased response frequency to capsaicin (**Fig. 8G**). These data are consistent with our *in vivo* findings **(Fig. 2**) and underscore the contributions of mEERL sEVs and CD8^+^ T cells to the transcriptome and function of tumor-infiltrating nociceptor neurons.

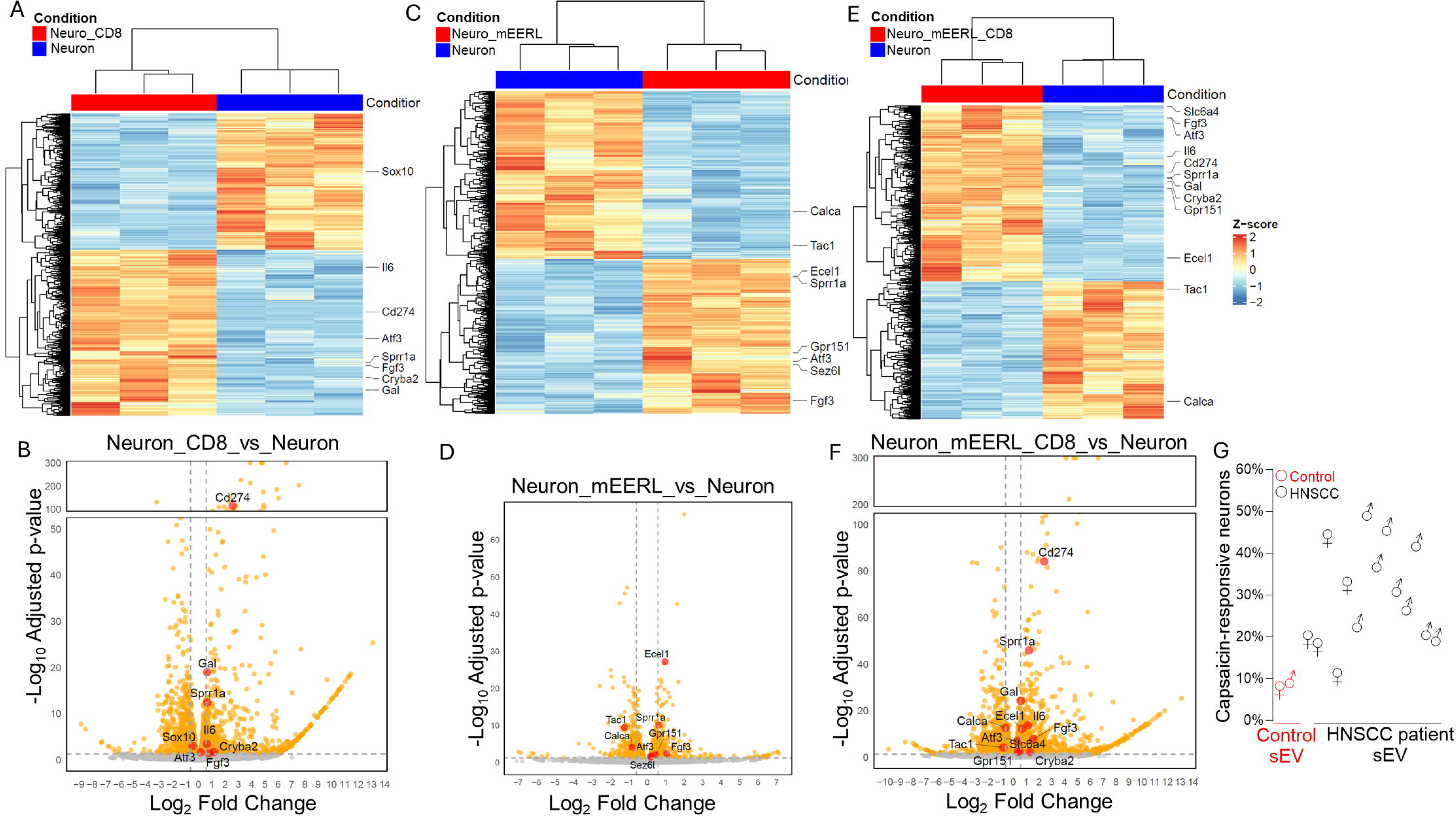

## Discussion

The infiltration of cancers by nociceptor neurons highlights the complex interactions that malignancies establish with their host, akin to the long-recognized connections of vascular and immune systems with tumors (*42–46*). In this study, we focused on the neural influences on disease progression, specifically in head and neck cancer and melanoma. Our findings demonstrate that tumor-released sEVs together with CD8^+^ T cells alter the transcriptome and sensitivity of tumor-infiltrating nociceptor neurons. Likely because of these changes, tumor-infiltrating nociceptor neurons alter the factors they release, contributing to a pro-tumorigenic tumor microenvironment.

Underlying these changes, we posit the existence of a feed-forward loop where tumor-released sEVs stimulate SP release from nociceptor neurons. This neuropeptide then triggers IL-6 release from mEERL cells. Concurrently, soluble factors released from mEERL cells enhance IL-6 production and release by nociceptor neurons. While our focus was primarily on IL-6, it is important to note that other factors (CCL2, CCL19, CXCL5, CD30L, CxCl16, TIMP1) were also induced by mEERL/nociceptor neuron co-cultures that likely also contribute to the tumor microenvironment’s immunosuppressive characteristics. These data matched the recent findings from Von Andrian and colleagues showing neuronal release of CXCL5 and CCL2 (*47*) as well as the one of Flavel presenting the release of IL-6 (*48*) For instance, one could imagine that the neuronal release of CCL2 might be important for retaining conventional dendritic cells within the tumors and the subsequent establishment of anti-tumor immunity.

Flow cytometric analysis reveals that this cytokine milieu recruits MDSCs to the tumor bed, a process significantly diminished in tumors implanted in nociceptor neuron-ablated animals. We demonstrated *in vitro* that this milieu not only stimulated MDSC differentiation and migration but also that factors released by nociceptors together with mEERL sEVs modulated the functionality of CD8^+^ T lymphocytes, enhancing their expression of immune checkpoint proteins PD-1, LAG3, and TIM3, while reducing the production of IFN-γ and IL-2. Moreover, interactions between mEERL sEVs and CD8^+^ T cells induce transcriptional changes on nociceptor neurons, leading them to express a neuronal injury-associated transcriptome. These findings collectively suggest that tumor cells, the neurons that infiltrate them, and the immune cells recruited to the site collaboratively create an immunosuppressive environment, thereby contributing to disease progression. This intricate interplay emphasizes the potential of targeting these neural and molecular interactions in therapeutic strategies to combat cancer.

These findings underscore the necessity of reassessing the impact of immunotherapies on patients with head and neck squamous cell carcinoma and melanoma, with consideration given to incorporating nerve-targeting therapies. Current strategies for developing new therapeutic targets in HNSCC are often inadequate which is partially due to the disease’s heterogeneity (*49, 50*). Although targeted therapies and immunotherapies represent significant advancements, their outcomes have been generally disappointing (*51–56*)). For example, Cetuximab, a leading targeted therapy for HNSCC, provides only modest clinical benefits with low response rates (*57, 58*). Similarly, PD-1 monoclonal antibody treatments like Pembrolizumab, though promising, show variable success largely dependent on the specific immune phenotype of the patient’s tumor (*59, 60*).

Given these challenges, it is crucial to explore how a tumor’s unique characteristics modulates the microenvironment as this will influence treatment responses. Our study has particularly focused on two pivotal immune cells, MDSCs and CD8^+^ T cells, which are integral to cancer progression and demonstrate the influence of nociceptor neurons and tumor sEVs on their number and functional status. Recent studies also highlight the significant role of the nervous system, including sensory neurons, in altering the tumor immune environment (*18, 19, 61–63*). Typically, neuro-immune interactions aim to maintain homeostasis, but tumor cells disrupt this balance, promoting immunosuppression (*64, 65*). Our findings are consistent with this. This maladaptive interaction suggests that targeting neurological pathways could complement existing immunotherapies. For instance, experiments in melanoma mouse models showed that silencing sensory neurons could enhance responses to anti-PD-L1 therapy, pointing to potential new directions in oncologic treatment strategies (*18*).

The future of cancer therapy may increasingly incorporate neuron-targeting strategies. This holistic view of the tumor microenvironment, which includes neural components, offers a broader perspective for developing more effective and personalized cancer treatment strategies. This could lead to innovative approaches that address not only the cancer cells but also the complex interplay of biological systems that support tumor growth and survival.

## Materials and Methods

### Cell lines

The mEERL (RRID:CVCL_B6J3), and Rab27^-/-^ (mEERL-Rab27a^-/-^/b^-/+^), cell lines have been characterized in previous studies (*26, 66, 67*). The mEERL and Rab27^-/-^ cells are cultured in E-medium, which consists of DMEM (Corning, cat# 10-017-CV) mixed with Ham’s F12 (Corning, cat#10-080-CV), supplemented with 10% fetal calf serum, 1% penicillin/streptomycin, 0.5 μg/ml hydrocortisone, 5 μg/ml transferrin, 5 μg/ml insulin, 1.36 ng/ml tri-iodothyronine, and 5 ng/ml epidermal growth factor (EGF). All cell lines are cultured at 37°C in an environment containing 5% CO_2_, and the culture medium is refreshed every three days.

### Study approvals

Animal studies were conducted within the controlled environments of the Sanford Research Animal Resource Center and a specific pathogen-free facility at Queen’s University. All procedures involving animals were conducted in accordance with the guidelines of the Canadian Council on Animal Care (CCAC) and the Queen’s University Animal Care Committee (UACC; 2023-2384).

Sanford Research has an Animal Welfare Assurance on file with the Office of Laboratory Animal Welfare (assurance number is A-4568-01). Sanford Health is a licensed research facility under the authority of the United States Department of Agriculture (USDA, certificate number 46-R-011). The Sanford Health Animal Research Program is accredited by AAALAC, Intl. All animal experiments conducted at Sanford Research were conducted under a Sanford Research approved IACUC protocol and all experimenters complied with ARRIVE guidelines.

### Animals

Mice were housed in individually ventilated cages with access to water and subjected to 12-hour light cycles; food was available ad libitum. C57BL6/J (Jax #000664), TRPV1^cre^ (Jax #017769) DTA^fl/fl^ (DTA; Jax #009669), were obtained from the Jackson Laboratory. As previously shown, (*18, 68–78*) animals were bred in-house to generate littermate control (TRPV1^wt^::DTA^fl/wt^), nociceptor reporter (Trpv1^cre^::td-tomato^fl/wt^) or ablated (*Trpv1^cre^::DTA^fl/wt^*) mice.

The Animal Resource Center (ARC) at Sanford Research is a specific pathogen-free facility. Mice are maintained in IVC Tecniplast Green line Seal Safe Plus cages. These cages are only opened under aseptic conditions in an animal transfer station. Aseptic technique is always used to change animal cages every other week; all cages have individual HEPA filtered air. Animal rooms are maintained at 75°F, 30-70% humidity, with a minimum of 15 air changes per h/cage. Rooms are maintained with a 14:10 light/dark cycle. Corncob bedding and nesting materials are autoclaved prior to use and are maintained in all cages. Animals were fed irradiated, sterile food (Envigo) and given acidified water (pH 2.8-3.0) *ad libitum*. There are a maximum of 5 mice/cage. Mice are observed daily by technicians. Abnormal behavior, signs of illness or distress, the availability of food and water and proper husbandry are monitored.

### Tumor implantation (mEERL and B16F10)

8-10-week-old C57BL/6 mice each weighed approximately 23 g at the start of experiments. The animals were uniquely identified by ear punches and cage numbers, and the investigators were blinded to group assignments when assessing the animals, such as during tumor measurements.

### Orthotopic (oral cavity) mEERL tumor implantation

Tumors were initiated into C57BL/6 mice as follows. Following anesthesia with ketamine (87.5 mg/kg)/xylazine (10mg/kg), each mouse was laid on its side. The mouth was gently opened, and the lower lip grasped with a pair of tweezers and pulled down to extend the tissue. A 23–25-gauge needle containing a suspension of mEERL cells was inserted into the crease of the mouse cheek, along the mandible. 1 x 10^5^ cells were slowly injected to orthotopically implant the cells in the submucosal space. Mice were placed under a heat lamp to recover. Once fully recovered, they were returned to their home cage. Mice were euthanized when tumor volume criteria were met, approximately 500 mm^3^. Tumors were measured every 7 days using calipers. Prior to tumor measurement, mice were anesthetized with isoflurane. Tumor volume was calculated using the following equation: (L x W^2^)/2. Following euthanasia, tumors were extracted and utilized for downstream assays.

### Cell lines

B16F10-mCherry-OVA (Matthew F. Krummel, UCSF), were cultured in complete Dulbecco’s Modified Eagle’s Medium high glucose (DMEM, Corning, #10-013-CV) supplemented with 10% fetal bovine serum (Seradigm, #3100) and 1% penicillin/streptomycin (Corning, #MT-3001-Cl), and maintained at 37°C in a humidified incubator with 5% CO_2_.

mEERL and mEERL Rab27^-/-^ cells were cultured as previously described (PMID: 30327461). Briefly, cells were cultured in DMEM (Corning, cat# 10-017-CV)/Ham’s F12 (Corning, cat# 10-080-CV), 10% sEV-depleted fetal calf serum, 1% penicillin/streptomycin, 0.5 µg/ml hydrocortisone, 5 μg/ml transferrin, 5 µg/ml insulin, 1.36 ng/ml tri-iodo-thyonine, and 5 ng/ml EGF. The cells were maintained at 37°C in a humidified incubator with 5% CO_2_.

All the cell lines tested negative for mycoplasma, and none are listed by the International Cell Line Authentication Committee registry (version 11). Non-commercial cell lines (B16F10-OVA) were authenticated using antibody (against OVA, eGFP, mCherry) and/or imaging as well as morphology and growth property. Commercial cell lines have not been further authenticated.

### Melanoma inoculation and volume measurement

Cancer cells were resuspended in Phosphate Buffered Saline (PBS, Corning #21040CV) and injected to the mice’s skin right flank (5×10^5^ cells; i.d., 100 μL). Growth was assessed daily using a handheld digital caliper and tumor volume was determined by the formula (L × W2 × 0.52). L = length and W = width.

### Trigeminal (TGM) Ganglia isolation

A midline incision was made on euthanized animals while in the prone position and this exposed the crown of the skull. A transverse cut was used to separate the brainstem from the spinal cord and the top of the skull was removed, thereby exposing the brainstem and TGM. The TGM ganglia were then harvested and utilized for downstream assays.

### Dorsal root ganglia isolation and co-culture

Dorsal root ganglia (DRG) were isolated from C57BL/6 or TRPV1^cre^::DTA^fl/wt^ mice following euthanasia. The mice were perfused with ice-cold HBSS and then underwent laminectomy to expose the spinal cord, after which the DRG were excised and immediately placed in ice-cold HBSS. The collected DRG were then embedded in 100 µl of CultureX (R&D Biosystems Cat# 3433-005-01) within a 30 mm cell culture dish. The CultureX was pipetted into the center of the dish and allowed to sit at room temperature for 5 min before the DRG were inserted into the matrix. Subsequently, the DRG were incubated at 37°C for 30 min, after which Ham’s F-12 medium supplemented with 10% FBS was added to the dish. After an overnight incubation, the DRG were either cultured alone or co-cultured with 3×10^4^ mEERL or Rab27^-/-^ cells. Three days later, conditioned media from these cultures were collected for downstream *ex vivo* experiments.

### Ex vivo generation of myeloid derived suppressor cells

Male C57BL/6 mice were euthanized, and bone marrow cells were isolated from their long bones following a previously described method (*79*). Briefly, the surrounding muscle tissue was grossly removed from the long bones, which were then soaked in serum-free RPMI for 5 min to ease the removal of any residual tissue. Subsequently, the bones were soaked in ethanol for another 5 min to ensure sterilization before being thoroughly rinsed in HBSS to remove any traces of ethanol. The epiphyseal ends of the bones were then cut open, and the marrow cells were flushed out by injecting RPMI into the marrow cavity using a syringe fitted with a 25-gauge needle. Once isolated, the bone marrow cells were cultured in RPMI supplemented with 10% fetal bovine serum (FBS). To generate myeloid-derived suppressor cells (MDSCs), the bone marrow cells were cultured in RPMI also containing 10% FBS, along with 40 ng/ml each of interleukin-6 (IL-6) and granulocyte-macrophage colony-stimulating factor (GM-CSF).

### Transwell migration assays

Bone marrow cells were harvested and stimulated to differentiate into MDSCs as described above. 2 x 10^4^ MDSCs were seeded onto a transwell (8 µm pore size, 3.0 µm polycarbonate membrane, Costar, catalogue #34028). In the well below, 1 mL of conditioned media from the designated co-cultures were added. Plates were kept in the incubator at 37°C for 12 h. The number of MDSCs that migrated onto the underside of the transwell were analyzed as follows. The top of the transwell was wiped with a Q-tip to remove all cells. The membrane was then fixed with ethanol and stained with crystal violet. The inserts were left to dry at room temperature for 15 min before the membrane was removed and mounted onto a glass slide and the underside (containing the migrated cells) analyzed by microscopy. ImageJ was utilized to quantify the number of cells. n=3 wells/group were analyzed and the experiment was repeated twice with similar results.

### Flow cytometry of bone marrow cells

Bone marrow cells (BMCs) were harvested as previously described and seeded onto 50 mm dishes each containing at least 2×10^5^ cells. These bone marrow cells were treated with condition media collected from mEERL cells, mEERL-Rab27^-/-^ cells, C57BL/6 DRG, TRPV1^cre^::DTA^fl/wt^ DRG, or their respective co-cultures (cancer cells and DRG). Positive controls were generated by treating bone marrow cells with IL-6 and GM-CSF as previously described. Negative controls were unstimulated bone marrow cells cultured in RPMI with 10% FBS. BMCs were incubated with the various condition media for up to 72 h after which they were collected, and viability was analyzed using trypan blue exclusion. BMCs were then washed, Fc-blocked, and stained with panel of fluorescent markers to identify specific immune cell populations (please see tables). Following staining, cells were resuspended in 200 uL FACS buffer and analyzed on a five-laser BD FACS Fortessa (BD Biosciences, San Jose, CA, USA). FCS files were exported, and compensation and analysis were conducted using FlowJo v9.7.

### Immunofluorescent staining

Formalin-fixed paraffin-embedded samples were cut at 5 µm thickness. Sections were deparaffinized through successive washes in: xylene (5 min), 100% ethanol (1 min), 90% ethanol (1min), 70% ethanol with 0.25% NH_3_ (1 hr), 50% ethanol (1min), water (1min). Slides were then incubated in heated antigen retrieval buffer (10 mM sodium citrate, 0.05% Tween 20, pH 6.0) for 30min and then rinsed in running water. Slides were then removed, tissues circled with Immedge pen and then incubated in blocking buffer (3% goat serum, 1% BSA, 1% cold fish skin gelatin, 0.1% Triton X-100, 0.05% tween 20, in 1x PBS, pH 7.2) for 30 min at room temperature. Following three washes in PBS, slides were incubated in Sudan Black for 30 sec followed by three PBS washes. Sections were incubated in primary antibody and maintained in a humidified chamber overnight at +4°C. The following day, slides were washed three times with PBS and incubated in secondary antibody, maintained in a humidified and dark chamber at room temperature for 1h. Slides were washed three times with PBS, incubated and coverslips were mounted using ProLong Glass antifade mounting medium (ThermoFisher, # P36980). Samples were analyzed by confocal microscopy (Nikon A1 TIRF).

### Antibodies used for immunofluorescent staining

Rabbit anti-mouse SP antibody (Thermo Fisher Scientific Cat# PA5-106934, RRID:AB_2854598) used at 1:1000; Rabbit anti-ATF3 antibody (Thermo Fisher Scientific Cat# PA5-106898, RRID:AB_2854562) used at 1: 500.

### Substance P ELISA

Condition media were collected from mEERL cells, DRG, or their co-culture. Co-cultures were generated as follows. Each co-culture contained 3 – 4 DRG which were extracted from C57BL/6 mice as previously described. DRG were placed onto Matrigel which was dropped on a 35 mm dish. DRG/Matrigel dishes were left undisturbed overnight in an incubator. The following day, 1×10^6^ mEERL cells were plate along the periphery of the Matrigel/DRG and the co-culture incubated for 48 hrs before collection. Condition media for single cultures was collected from mEERL cells when plates were 80% confluent and from DRG following 48 – 72h after being plated on Matrigel. The concentration of SP in harvested condition media was estimated using a standard curve with a SP EIA kit from RayBiotech ( catalog# EIA-SP).

### IL-6 ELISA

Condition media from mEERL cells alone, DRG alone, or their co-culture were harvested as described for the SP ELISA. For determination of SP mediated IL-6 release, mEERL cells were plated on 35 mm dishes in serum-free media overnight. The following day, 50 nM substance P (Sigma Aldrich, acetate salt hydrate, Cat#S6883) alone, or with 100 uM NK1R antagonist (Tocris Bioscience, L-732,138) was added. Cells were incubated with treatment for 48h prior to conducting IL-6 ELISA which was performed as per manufacturer directions (RD Systems, cat # M6000B).

### Cytokine array

Cytokine arrays were purchased from RayBiotech (catalog # AAM-CYT-3). Condition media were harvested from *in vitro* cultures of DRG alone, mEERL cells alone, or culture of mEERL cells with DRG as described above. Cytokine arrays were processed per manufacturer’s recommendations. Briefly, arrays were blocked at room temperature and then treated with undiluted condition media overnight at 4°C. The following day, arrays were treated with biotinylated antibody cocktail for 2 h at room temperature, and then incubated for 2 h with HRP-Streptavidin diluted in blocking buffer. Arrays were treated with detection buffer and imaged on a Li-COR Odyssey imaging system.

### Antibodies used for flow cytometry

The antibodies used for flow cytometry are listed in the table. A live/dead stain (Invitrogen, Fixable blue cat. # L34962) was also used.

### Table A

Table A:
| Antibody | Vendor | Catalogue # | RRID |
| --- | --- | --- | --- |
| Anti-CD8 | BioLegend | 100712 | AB_312751 |
| Anti-PD-1-PE-Cy7 | BioLegend | 109110 | AB_572017 |
| Anti-LAG3-PE | BioLegend | 125208 | AB_2133343 |
| Anti-TIM3-PerCP/Cyanine5.5 | BioLegend | 134012 | AB_2632736 |
| IFNγ-FITC | BioLegend | 505806 | AB_315400 |
| TNF-BV510 | BioLegend | 506339 | AB_2563127 |
| IL-2-Pacific Blue | BioLegend | 503820 | AB_2127161 |
| CD45.2-AF488 | BioLegend | 109815 | AB_492869 |
| CD3-BV510 | BioLegend | 100233 | AB_2561387 |
| CD4-BV421 | BioLegend | 100543 | AB_10898318 |
| MHCII-APC | eBioscience | 17-5321-82 | AB_469455 |
| NKp46-PECy7 | BioLegend | 137617 | AB_11218594 |
| CD11b-PE | eBioscience | 12-0112-81 | AB_465546 |
| CD8a-APCCy7 | BioLegend | 100713 | AB_312752 |
| CD11c-BV785 | BioLegend | 117335 | AB_11219204 |
| CD25-PerCPcy5.5 | eBioscience | 45-0251-80 | AB_914323 |
| GR-1-AF700 | BioLegend | 108421 | AB_493728 |
| F4v/80-BV605 | BioLegend | 123133 | AB_2562305 |
| CD45.2-BV605 | BioLegend | 109841 | AB_2563485 |
| MHCII-PE-Cy7 | BioLegend | 107629 | AB_2290801 |
| CD11b-PE-Cy5 | BioLegend | 101210 | AB_312793 |
| Ly6G-AF700 | BioLegend | 127622 | AB_10643269 |
| Ly6C-FITC | BioLegend | 128005 | AB_1186134 |
| CD3-PE | BioLegend | 100206 | AB_312663 |
| CD4-APC | eBioscience | 17-0042-82 | AB_469323 |
| CD8a-BV421 | BioLegend | 100753 | AB_2562558 |
| CD335 NKp46-BV510 | BioLegend | 137623 | AB_2563290 |

### Tumor dissociation and flow cytometry

*In vivo* studies utilized 8-week-old male C57BL/6 or TRPV1^cre^::DTA^fl/wt^ mice. For the experimental procedure, 100,000 mEERL cells were orthotopically injected into the oral cavity, following previously established methods and as described above (*10, 80*). Twenty-five days after the injection, the mice were euthanized, and the tumors were harvested. The tumors were then processed according to the MACS Tumor Dissociation protocol to ensure a single-cell suspension suitable for subsequent analyses. Cell viability post-dissociation was assessed using TO-PRO-3 staining (Thermo Fisher, cat# T3605), and viable cells were sorted using an Accuri flow cytometer. For phenotypic analysis of the tumor-infiltrating cells, 1×10^6^ cells were stained with one of three fluorescent antibody panels. The panels used included a 12-color panel, a 9-color panel specifically designed for Ly6C/G identification, and a 4-color panel, detailed in accompanying tables. Flow cytometry data files were then compensated and analyzed using FlowJo software, providing detailed insights into the cellular composition and immune phenotype of the tumor microenvironment. This comprehensive approach allows for a robust assessment of the impact of tumor and immune cell interactions within the tumor microenvironment.

### Co-culture of sEVs and DRG neurons

DRG were extracted from male C57BL/6 mice and placed into complete DMEM medium (Corning, 10-013-CV) supplemented with 50 U/ml penicillin, 50 µg/ml streptomycin (Corning, MT-3001-Cl), and 10% FBS (Seradigm, 3100). The neurons were then dissociated using phosphate-buffered saline (Corning, 21-040-CV) enriched with 1 mg/ml collagenase IV (Sigma, C0130) and 2.4 U/ml dispase II (Sigma, 04942078001). This mixture was incubated for 80 min at 37°C. Afterward, the ganglia were triturated using glass Pasteur pipettes of decreasing sizes in complete DMEM, followed by centrifugation over a 10% BSA gradient. The cells were then plated on cell culture dishes coated with laminin (Sigma, L2020).

The plated cells were cultured in DMEM medium (Gibco, 21103-049) completed with 10% FBS, 1% penicillin–streptomycin, non-essential amino acids (Corning, 25-025-Cl), and additional supplements including β-mercaptoethanol (Gibco, 21985-023), L-glutamine (VWR, 02-0131), 0.05 ng/µl sodium pyruvate (Corning, 25-000-Cl), a specified concentration of NGF (Life Technologies, 13257-019), and 0.002 ng/µl GDNF (PeproTech, 450-51-10). After 2h, the cells were co-cultured either with small extracellular vesicles (sEVs; 3 µg in 200 µl) or PBS, both in the presence of a peptidase inhibitor (1 µM). Conditioned media were collected after 48h for further analysis.

### Isolation of sEVs

sEVs from mEERL cells were isolated by differential ultracentrifugation as previously described (*26*). Briefly, conditioned media from mEERL cells was collected and spun in a Thermo Legend X1R centrifuge at 300 *x g* for 10 min. The supernatant was collected and spun at 2000 *x g* for 10 min. The supernatant was collected and spun in a Sorval RC6 centrifuge at 10,000 *x g* for 30 min. The resulting supernatant was collected and spun in a Sorval WX80 Ultracentrifuge at 110,000 *x g* for 2 h. The resulting pellet contains the sEVs and was washed with sterile PBS and spun again (110,000 x g, 2 hrs). The supernatant was discarded and the pellet (sEVs) resuspened in 200 µl of sterile PBS, aliquoted and stored at -80 °C until used. sEVs were validated by Nanosight particle analysis.

### Co-culture of CD8^+^ T cells and DRG neurons condition media

Spleens were harvested from naïve male mice into cold PBS supplemented with 5% FBS and kept on ice. The tissues were mechanically dissociated and then passed through a 70 µm strainer. Red blood cells were lysed using RBC lysis buffer (Life Technologies, A1049201) for 2 min, and the remaining cells were counted using a hemocytometer. Splenocytes were isolated via magnetic sorting using a specific kit (Stem Cell, 19853A) and subsequently cultured in DMEM supplemented with 10% FBS, 1% penicillin–streptomycin, non-essential amino acids (Corning, 25-025-Cl), β-mercaptoethanol (Gibco, 21985-023), L-glutamine (VWR, 02-0131), and sodium pyruvate (Corning, 25-000-Cl).

The splenocytes were then stimulated under Tc1 conditions using 2 µg/ml of anti-CD3 and anti-CD28 antibodies (Bio X Cell, BE00011, BE00151), 10 ng/ml recombinant IL-12 (BioLegend, 577008), and 10 µg/ml anti-IL-4 antibody (Bio X Cell, BE0045), in a 96-well plate. After 48 h of stimulation, the cells were transferred to uncoated plates and exposed to either purified sEVs or conditioned media from sEV/DRG neuron co-cultures for an additional 72 h.

Subsequently, the expression of checkpoint proteins PD-1, LAG3, and TIM3, as well as the secretion of cytokines IFNγ, TNFα, and IL-2, were analyzed using flow cytometry, specifically using an LSRFortessa or a FACSCanto II (Becton Dickinson). The quantification of cytokine expression was performed after *in vitro* stimulation, providing detailed insights into the functional status of the CD8^+^ T cells in response to the experimental treatments. This comprehensive approach facilitates a deeper understanding of how CD8^+^ T cell functionality can be modulated by external factors such as sEVs and DRG neuron-derived signals within the immune microenvironment.

### Intracellular cytokine staining

Cytotoxic CD8^+^ T cells were stimulated with phorbol-12-myristate 13-acetate (PMA; 50 ng/ml, Sigma-Aldrich, P1585), ionomycin (1 µg/ml, Sigma-Aldrich, I3909), and Golgi Stop (1:100, BD Biosciences, 554724) for 3 h to activate them and halt protein transport, enabling cytokine accumulation. After stimulation, the cells were washed with FACS buffer, which consists of PBS supplemented with 2% fetal calf serum and EDTA. This was followed by staining the cells with Viability Dye eFluor 780 (eBioscience, 65-0865-14) for 15 min at 4°C to assess cell viability.

Post viability staining, the cells underwent another round of washing and were then stained for 30 min at 4°C with several antibodies: anti-CD8-APC (BioLegend, 100712), anti-PD-1-PE-Cy7 (BioLegend, 109110), anti-LAG3–PE (BioLegend, 125208), and anti-TIM3-PerCP/Cyanine5.5 (BioLegend, 134012). These stains were used to identify the CD8^+^ T cells and to evaluate their expression of various immune checkpoint proteins.

Following surface staining, the cells were fixed and permeabilized using a kit (BD Biosciences, 554714) to allow for intracellular staining. The cells were then stained for IFNγ–FITC (BioLegend, 505806), TNF–BV510 (BioLegend, 506339), and IL-2–Pacific Blue (BioLegend, 503820) to detect the production of key cytokines that indicate cellular activation and function. The final analysis of the stained cells was conducted using flow cytometry, employing either a LSRFortessa or FACSCanto II system (Becton Dickinson), providing detailed insights into the functional status and health of the CD8^+^ T cells in response to stimulation. This multi-parameter flow cytometry approach is essential for understanding the immune functionality and regulation of cytotoxic T cells under various conditions.

### RNA sequencing of triple co-cultures and data processing

Naive TRPV1^cre^::tdTomato^fl/WT^ DRG neurons, specifically 4 × 10^4^ in number, were co-cultured (1:10 ratio) with mEERL-derived small extracellular vesicles (sEVs), CD8^+^ T cells, or a combination of both, each within T cell medium.The medium was supplemented with neurotrophic factors: 0.05 ng/µl neuron growth factor (NGF) from Life Technologies (Cat# 13257019) and 0.002 ng/µl glial cell line-derived neurotrophic factor (GDNF) from PeproTech (Cat# 450-51-10), to support neuronal survival and function.

After 48 h of co-culturing, the cells were collected, and the TRPV1-expressing neurons, identifiable by their tdTomato fluorescence, were purified using a FACSAria IIu cell sorter (Becton Dickinson). This sorting process ensures that subsequent analyses or experiments are conducted on a homogeneous population of TRPV1^+^ neurons, eliminating any non-neuronal or non-TRPV1-expressing cells that could confound results. This methodical approach facilitates the study of specific interactions between DRG neurons and immune cells or factors within the controlled conditions of an *in vitro* system.

RNA-sequencing libraries of TRPV1 neurons were constructed using the Illumina TruSeq Stranded RNA LT Kit, adhering closely to the manufacturer’s instructions provided by Illumina. Sequencing of these libraries was carried out at Fulgent Genetics. The sequencing reads were then aligned to the Mouse mm10 reference genome (GenBank assembly accession GCA_000001635.2) using the STAR software version 2.7. After alignment, reads that mapped to genic regions were quantified using the featureCounts function from the subread package version 1.6.4.

Gene expression levels across the samples were quantified in terms of Transcripts Per Million (TPM), which facilitates comparison between samples by normalizing for both sequencing depth and gene length. Hierarchical clustering of gene expression data was performed using the heatmap.2 function from the gplots package in R (version 3.1.3), employing the ward.D2 method to discern patterns and relationships in gene expression among the samples. For differential gene expression analysis, DeSeq2 version 1.28.1 was utilized to identify genes that were significantly upregulated or downregulated under different experimental conditions. The results of these analyses, including all relevant data, have been deposited in the NCBI’s Gene Expression Omnibus (GEO), accessible under the accession number GSE205864. This comprehensive approach provides a robust framework for understanding the transcriptional changes in TRPV1 neurons in response to various experimental treatments.

### Quantitative polymerase chain reaction for MDSC genes

Real-time quantitative reverse transcription (RT-qPCR) was conducted to analyze the levels of MDSC-associated transcripts in the generated MDSC populations. The process began with the extraction of total RNA using Qiazol extraction reagent, followed by further purification through phenol-chloroform extraction protocols. The quality and quantity of the extracted RNA were assessed using a NanoDrop spectrophotometer. Subsequently, cDNAs were synthesized from the RNA samples using a High-Capacity cDNA Reverse Transcription kit from Applied Biosciences. Gene expression analysis was performed using real-time quantitative RT-PCR on a CFX-96 system from BioRad. Specific primers used for amplifying the gene products are detailed above. The mRNA levels of the genes of interest were quantified using the comparative threshold cycle (Ct) method. This involves normalizing the expression level of each gene of interest to that of a housekeeping gene, β-actin, to account for variations in RNA input and efficiency of the RT reaction across different samples. This normalization is crucial for accurate, reproducible, and meaningful quantification of gene expression, facilitating the comparison of mRNA levels across different experimental conditions and samples.

### Statistical analysis

GraphPad Prism (version 10.0.3, 2023) was used for all statistical analyses. The specific statistical tests for each experiment are noted below in the corresponding figure legend.

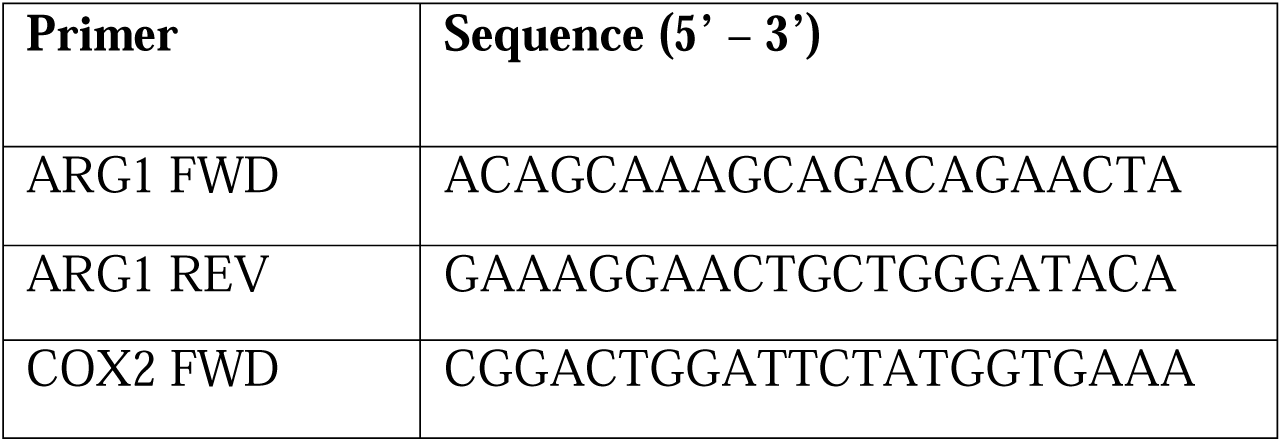
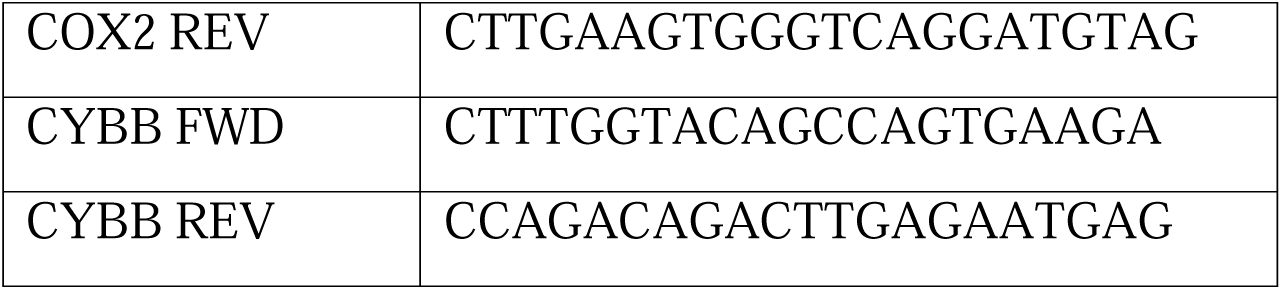

### Tumor growth curves

Two-way ANOVA with post-hoc Tukey test or two-sided unpaired Student’s t-test.

### Gene/protein expression differences

Student’s t-test or one-way ANOVA with post-hoc Tukey test.

### ELISA

One-way ANOVA with post-hoc Tukey test.

### Flow cytometry data

Two-sided unpaired Student’s t-test or one-way ANOVA with post-hoc Tukey test.

### MDSC migration assay

One-way ANOVA with post-hoc Tukey test.

## Acknowledgments

We thank the Flow Cytometry Core (Sanford Research, supported by National Institute of General Medical Sciences, Center of Biomedical Research Excellence P30GM145398) for their services and expertise towards this project.

## Author contributions

Conceptualization: PDV, ST, ACR
Methodology: ACR, MA, AW,
Investigation: ACR, MA, AW
Visualization: TE, ARN, MB, ACR
Supervision: PDV, ST
Writing—original draft: PDV
Writing—review & editing: PDV, ST, ACR, MRN, MB, TE
Project administration: PDV, ST
Funding acquisition: PDV, ST
Formal analysis: TE, ARN

## Data and materials availability

All data are available in the main text or the supplementary materials. Upon request, cell lines will be made available following a materials transfer agreement (MTA). The RNA sequencing dataset has been deposited in the NCBI’s Gene Expression Omnibus (GEO), accessible under the accession number GSE205864.

