## Supplementary figures and images for "TUMOR-INFILTRATING NOCICEPTOR NEURONS PROMOTE IMMUNOSUPPRESSION"

### Supplemental Figure 1

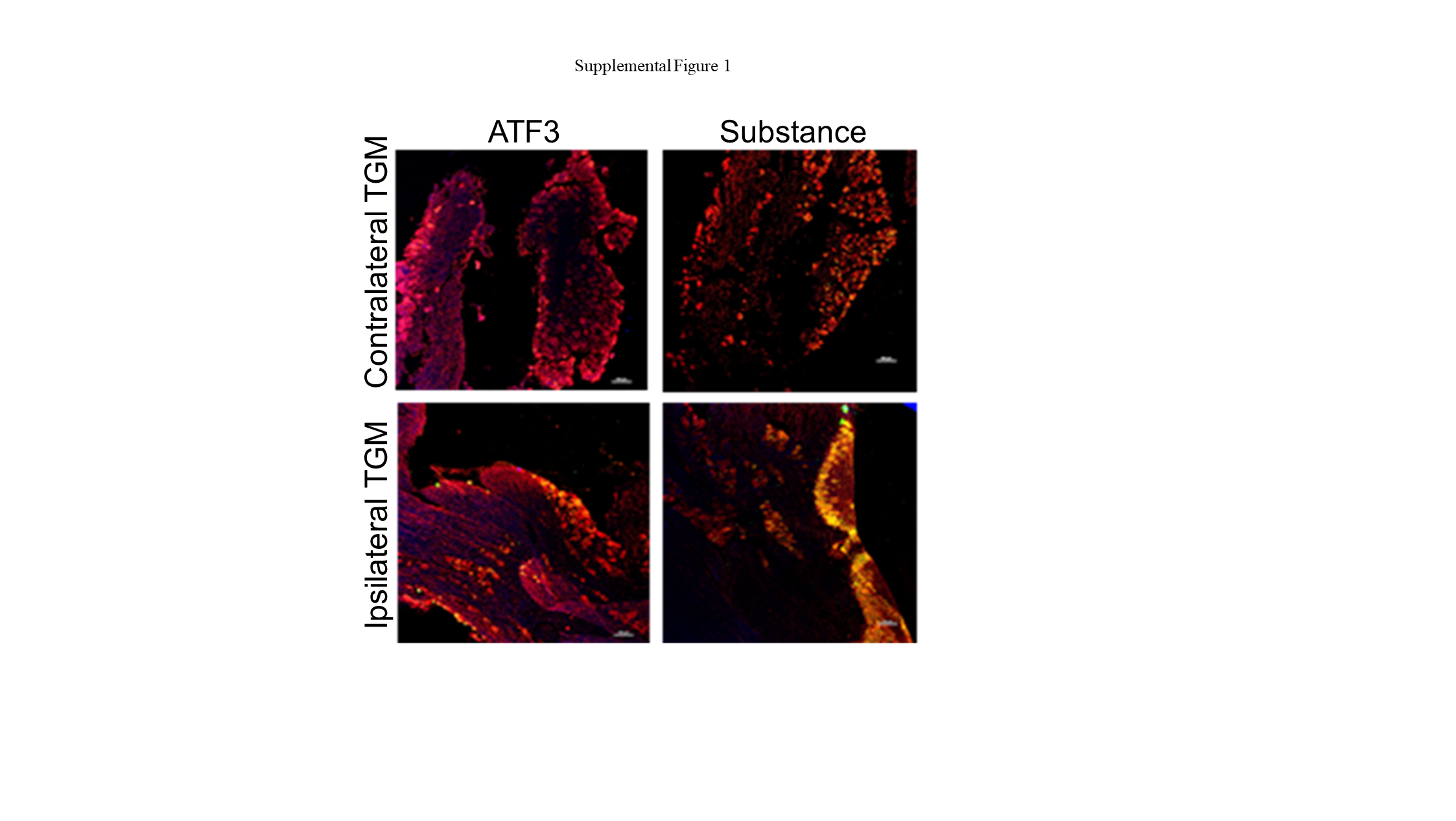

### Supplemental Figure 2

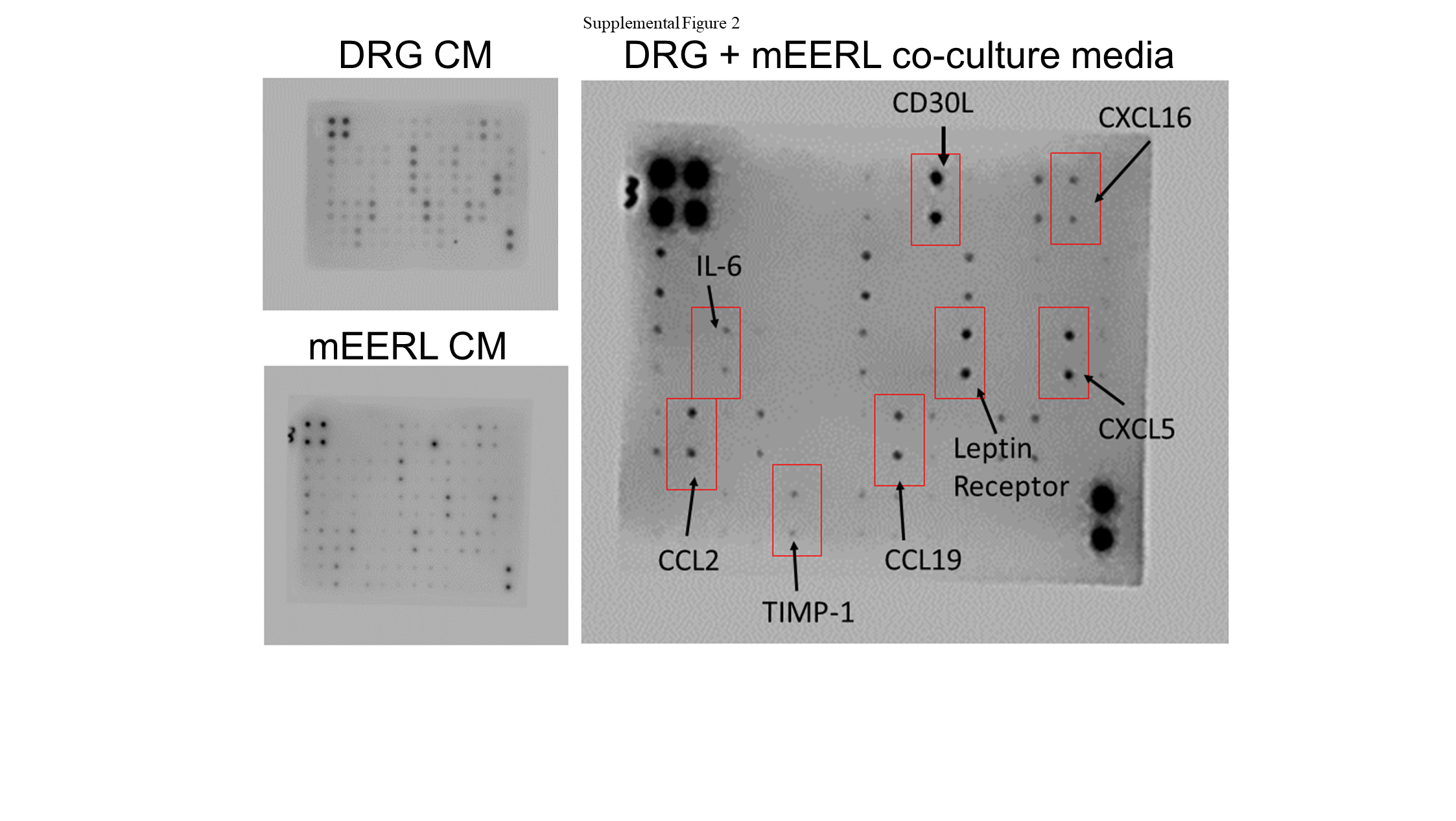

### Supplemental Figure 3

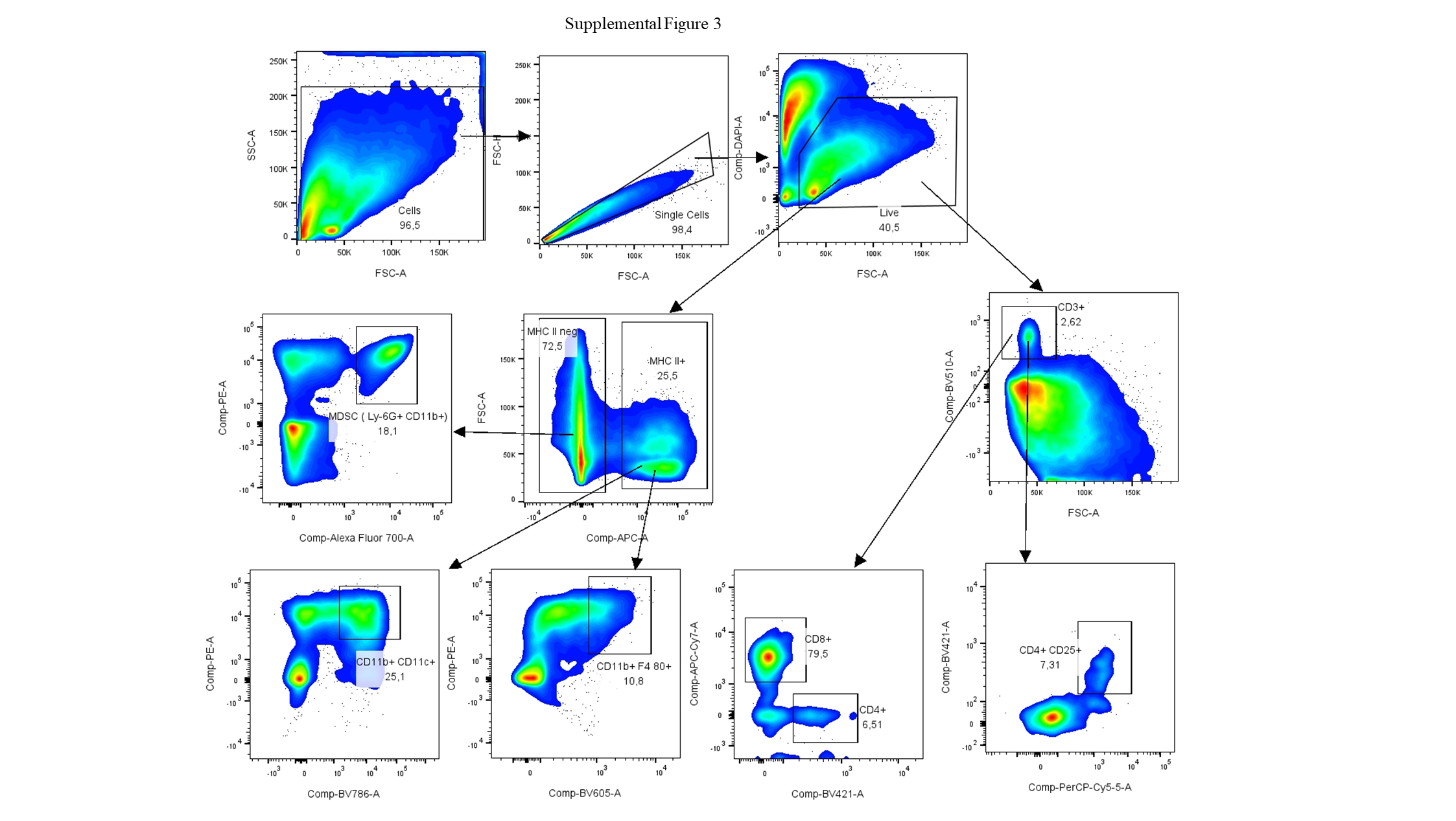

### Supplemental Figure 4

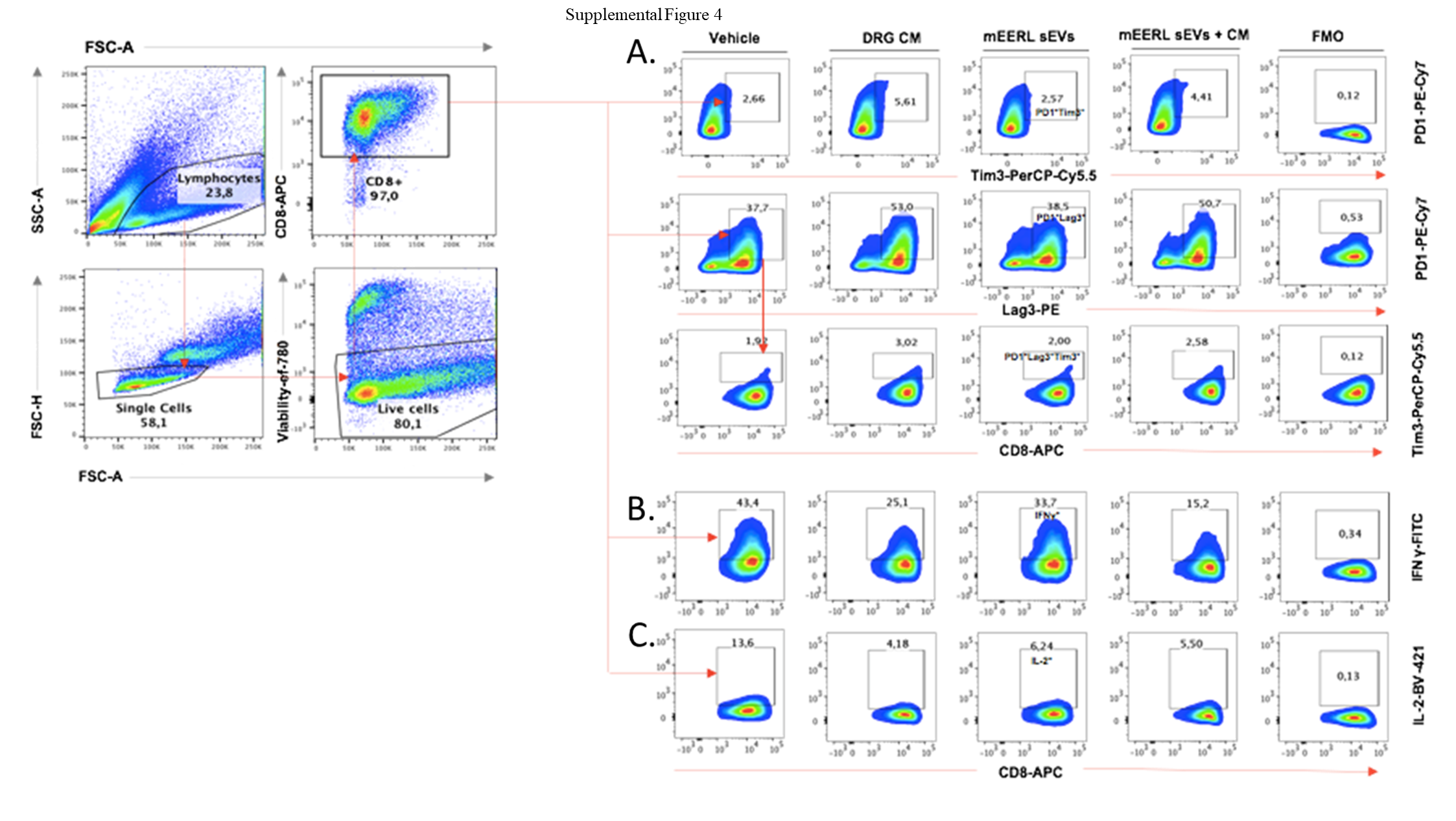
